# Analysis of iPLA2-VIA Gene Expression in The CNS of Larvae and Adult *Drosophila Melanogaster* Flies

**DOI:** 10.1101/2024.07.17.603941

**Authors:** Gabriel Moura

## Abstract

This study aimed to investigate the expression levels of the iPLA2-VIA gene (FlyBase: FBgn0035800) in the nervous tissue of *Drosophila melanogaster* (FlyBase: FBsp00000008) larvae and adults. The gene was identified using the FlyBase database and the HGNC database. The results of our experiments indicated that the expression levels of the gene were consistent between the larval and adult stages, with higher expression of Larvae than of Adult. We also found that the gene was expressed in the CNS, digestive system, fat body, salivary glands, ovaries, and accessory glands. Furthermore, PLA2G6 (Gene ID: 8398) knockout resulted in lethality, fertility problems, tissue formation defects, and neurological abnormalities, highlighting its essential biological roles in neural health and disease. Our findings could inform future studies focused on the gene’s involvement in neural development and its potential as a therapeutic target for neurodegenerative diseases.

## 2 Introduction

The *Drosophila melanogaster* model system is widely utilized in genetic research due to its genetic tractability, short lifecycle, and conserved biological processes with higher eukaryotes (Staats *et al*, 2018) One gene of interest within this system is the iPLA2-VIA gene, a homolog of the human PLA2G6 gene. The iPLA2-VIA gene encodes enzymes involved in lipid metabolism, which play a critical role in membrane remodeling, signal transduction, and energy metabolism (Balsinde, 2005). These enzymes are crucial for maintaining cellular homeostasis and function, particularly in the nervous system. Mutations in the human PLA2G6 gene have been associated with a spectrum of neurodegenerative disorders, including infantile neuroaxonal dystrophy (INAD), neurodegeneration with brain iron accumulation (NBIA), and Parkinson’s disease (Bohlega *et al*, 2016) (Fig 1). These conditions underscore the importance of iPLA2-VIA/PLA2G6 in neural health and disease, making it imperative to understand the function and regulation of the gene in model organisms like Drosophila. Given the significant role of iPLA2-VIA in neural integrity and the involvement of PLA2G6 in neurodegenerative diseases (Lin *et al*, 2018), we aimed to investigate the expression of the iPLA2-VIA gene specifically in the nervous tissue of *Drosophila* larvae and adults. Our study involved two main experimental groups: larval nervous tissue (control group) and adult nervous tissue. This approach allowed us to compare the gene expression patterns in different developmental stages and to obtain information on the temporal regulation of iPLA2-VIA. Since understanding the expression patterns of iPLA2VIA in *Drosophila* can provide valuable information on the role of the gene in neural development and function. Therefore, this research may contribute to the broader understanding of lipid metabolism in neurobiology and offers potential implications for studying human neurodegenerative diseases. By comparing gene expression in larval and adult nervous tissues, our objective is to uncover developmental dynamics that may be relevant to both basic and applied neurogenetic research. We hypothesized that the expression of iPLA2-VIA will exhibit significant variation between larval and adult nervous tissues, reflecting the gene’s involvement in the developmental processes and functional requirements of the nervous system at different life stages.

**Fig. 1.**
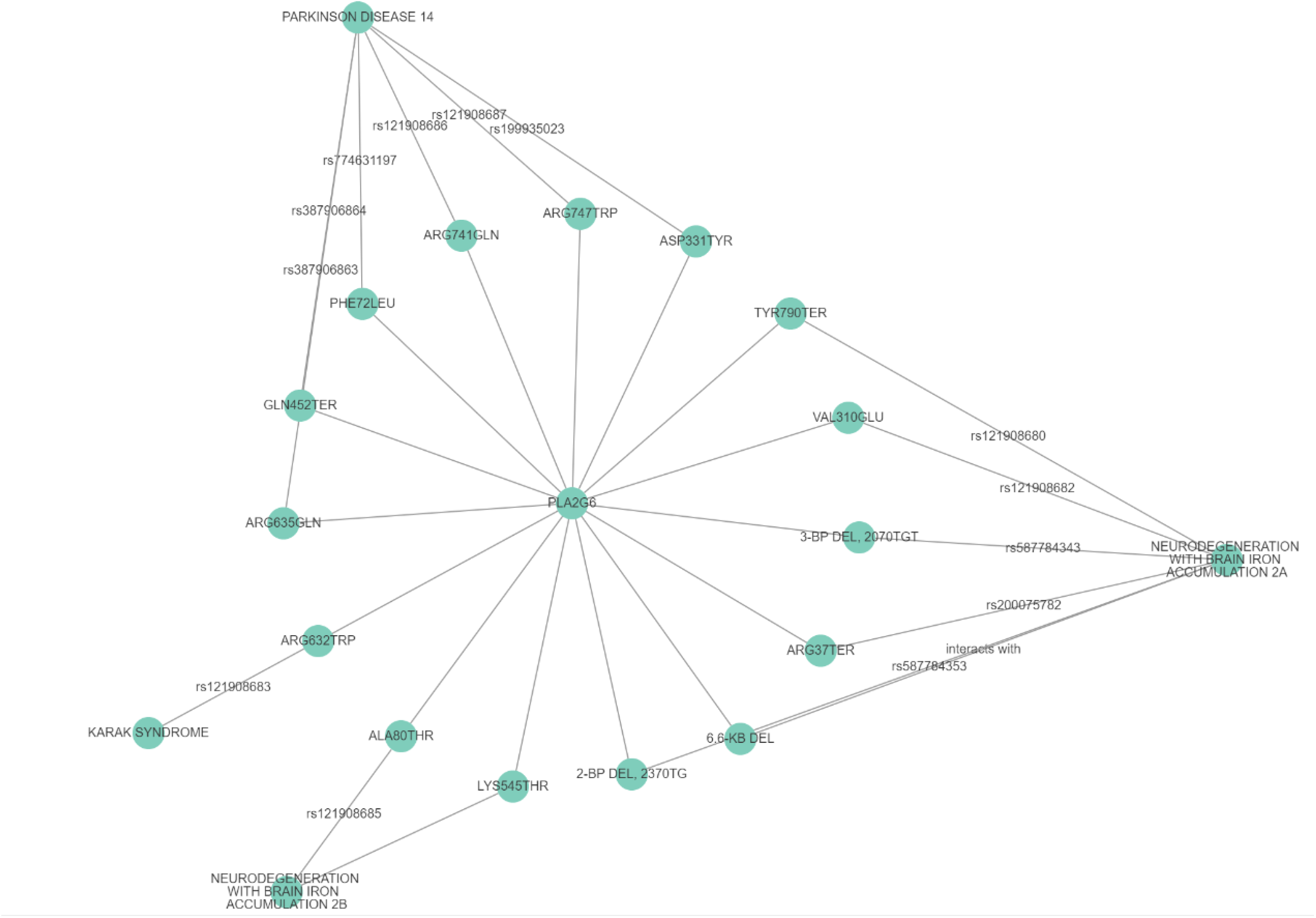
Reported Network of Alleles with PLA2G6 curated by OMIM with their respective Mutation, dbSNP, and Phenotype/Disease Expression.

## 3 Methods

We began our study by utilizing the FlyBase database to gather comprehensive information about the iPLA2-VIA gene in *Drosophila melanogaster*. The gene was identified as a protein-coding gene encoding two types of enzymes: phospholipase A2 (EC 3.1.1.4) and ADP-dependent medium-chain-acyl-CoA hydrolase (EC 3.1.2.19). We employed the “Jump to Gene (J2G)” search tool to locate iPLA2-VIA.

We looked at the CNS expression levels of the gene using the modENCODE and FlyAtlas2 transcriptomes, and used the MARRVEL link to identify the closest human ortholog, PLA2G6. Through the HGNC database, we verified that PLA2G6 has three predicted transcripts and is expressed in multiple human tissues, including the kidney, brain, and liver. Based on this, we began by accessing the cDNA sequence under the “sequence” tab. This sequence, which is more specific in detailing the base pairs (bp) for each alternating codon, facilitated easier primer design. We then explored single nucleotide variants (SNVs) within the sequence, identifying one and determining it did not alter the protein-coding sequence as it was in the untranslated region (UTR). Using the European Variation Archive (EVA), we assessed the prevalence of this variant in the *Drosophila* Genetic Reference Panel (DGRP), finding it to be uncommon with a 23% incidence rate. We repeated this process for another variant and found it more common with an incidence rate of 60%, probably due to the higher abundance of adenine base pairs in the DNA sequence.

For primer design, we used the Integrated DNA Technologies (IDT) website. After navigating to the PrimerQuest Tool, we manually entered the template sequence from the iPLA2-VIA Ensembl page and selected “PCR: 2 primers”. The software produced five sets of primers, different in their starting and stopping positions and GC content, where only one was chosen Table 1. We customized the assay design using data from Ensembl to meet our experimental criteria, and once the primers were designed accordingly, we added the forward and reverse primers to the order. The primers were named by “sequence name,” and standard concentration and purification parameters were used.

**Table 1.**
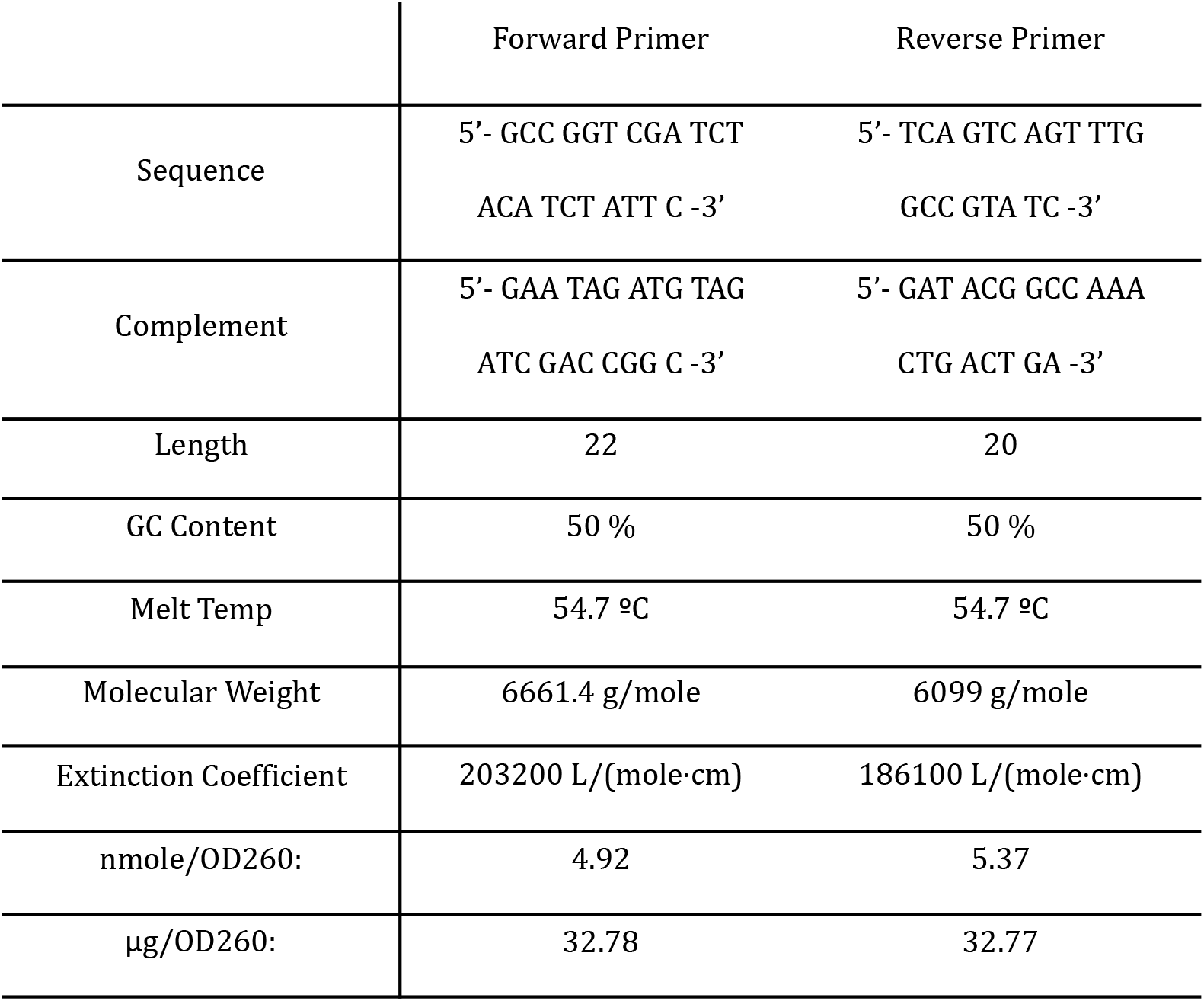
Primers Set.

### 3.1 Sample Preparation and RNA Collection

We began by collecting the nervous tissues from both larvae and adult *Drosophila* that were pestled in a test tube. These tissues were then homogenized in Trizol reagent to ensure efficient cell lysis and preservation of RNA integrity. The homogenate was incubated at room temperature for 5 minutes to allow the complete dissociation of nucleoprotein complexes. We then added chloroform to the mixture, vigorously shook it for 15 seconds, and let it stand at room temperature for another 2-3 minutes. This step was crucial in separating the mixture into aqueous and organic phases. We centrifuged the sample at 12,000 × g for 15 minutes at 4°C, which resulted in the separation of RNA (in the aqueous phase) from DNA and proteins. The aqueous phase, which contained the RNA, was carefully transferred to a new tube. We added 250*μ*L of 100 percent isopropanol to precipitate the RNA. After a 10-minute incubation at room temperature, the mixture was centrifuged at 12,000 × g for 10 minutes at 4°C. This step resulted in the formation of a visible RNA pellet. To purify the RNA, the pellet was washed with 1 mL of 75 percent ethanol, briefly vortexed, and centrifuged at 7500 × g for 5 minutes at 4°C. The ethanol was then carefully removed, and the pellet was allowed to air-dry for about 10 minutes to remove any residual ethanol. The dried RNA pellet was resuspended in 20*μ*L of RNase-free water. To ensure the RNA was fully dissolved, we incubated the tube at 55-60°C for 10-15 minutes. The resulting RNA solution was then stored at -70°C until further use. We measured the concentration and purity of the isolated RNA using a Nanodrop spectrophotometer. This step was essential for ensuring that the RNA samples were suitable for subsequent cDNA synthesis. Each sample’s RNA concentration was recorded to standardize the input for cDNA synthesis.

We synthesized cDNA from 2*μ*g of RNA for each sample by mixing 2*μ*L RNA, 2*μ*L oligo-dT primer, 2*μ*L random hexamer primers, 2*μ*L dNTPs, and RNase-free water to a total volume of 20*μ*L. Then the reaction mixture was incubated at room temperature for 10 minutes, followed by incubation at 50°C for 50 minutes. The reaction was stopped by heating at 85°C for 5 minutes. After a brief centrifugation, the cDNA was stored at -20°C.

### 3.2 Primer Design and PCR Amplification

We designed primers for the PCR amplification of iPLA2-VIA cDNA. The primers, synthesized by IDT, were rehydrated to a stock concentration of 100*μ*M and diluted to a working concentration of 10*μ*M. The lyophilized primers were rehydrated with RNase-free water at a concentration of 100*μ*M and mixed well by pipetting. Each PCR reaction was prepared in a fresh PCR tube with 2*μ*L template cDNA, 2.5*μ*L forward primer (10*μ*M), 2.5*μ*L reverse primer (10*μ*M), 25*μ*L PCR master mix (containing polymerase, dNTPs, Magnesium, and 18*μ*L RNase-free water, resulting in a final volume of 50*μ*L per reaction. The Thermocycling Conditions for the PCR were performed by initial denaturation at 98 ° C for 30 seconds followed by 25 cycles of:

|  |
| --- |
| Denaturation at 98 ° C for 10 seconds |
| Annealing at 60°C for 10 seconds |
| Extension at 72°C for 30 seconds |
| Final extension at 72 ° C for 5 minutes |

To analyze the PCR products, we performed agarose gel electrophoresis. We prepared a 0.75% agarose gel by mixing 0.45 g of agarose with 60 mL of 1x TAE buffer in an Erlenmeyer flask. The mixture was microwaved until the agarose dissolved completely. After cooling for 10 minutes, the solution was poured into a gel casting tray with a comb to form wells. Once the gel solidified, 2*μ*L of blue loading dye was added to 10*μ*L of each PCR sample. The samples were then loaded into the wells alongside a molecular weight marker. The gel was run at 100 V for 45 minutes to allow the DNA fragments to migrate. After running the gel, it was stained with ethidium bromide for 5 minutes and imaged using a Bio-Rad GelDoc imager under UV light.

## 4 Results

Our PCR and gel electrophoresis yielded visible bands under UV light ; however, it was far away from the expected position. Despite meticulously following the protocol, contamination or technical errors could have led to the absence of detectable PCR products. The data of the expression pattern of modENCODE and FlyAtlas2 indicated expression in the CNS, digestive system, fat body, salivary glands, ovaries and accessory glands, and a knockout of iPLA2-VIA resulted in lethality, fertility problems, tissue formation defects, and neurological abnormalities, highlighting its essential biological roles.

We expected for the Larvae to have higher expression of this gene compared to the Adult, due to the complexity of gene regulation in early form of an individual Based on the BLAST results, the top hit for the forward primer is within the range from 9861444 to 9861465, and for the reverse primer is within the range from 9861022 to 9861036 in the *Drosophila melanogaster* genome.

We can calculate the expected size of the PCR product by subtracting the starting position of the forward primer from the ending position of the reverse primer. Therefore, the expected size = (9861465 - 9861022) + 1 = 444 base pairs. The larvae had a higher concentration of RNA than the Adult, as you can see by the difference in opacity.

## 5 Discussion

Our study aimed to investigate the expression levels of the iPLA2-VIA gene in the nervous tissue of *Drosophila* larvae and adults. This investigation was motivated by the known roles of the gene in lipid metabolism and its association with neurodegenerative diseases in humans. The research involved isolating RNA from nervous tissues, performing RT-PCR, and analyzing the resulting expression levels and patterns. The anticipated outcomes were the identification of distinct expression levels of iPLA2-VIA between the larval and adult stages, providing insights into its developmental regulation and potential functional roles. The results of our experiments indicated that iPLA2-VIA is expressed in both larval and adult nervous tissues. We anticipated higher expression levels in larvae due to the increased complexity and functional demands to form a mature nervous system, which was confirmed. Since the concentration of iPLA2-VIA in larvae was greater than the adult, this could indicate that a mutation in the gene in the early development might lead to future neurological diseases in flies. Banerjee et al. 2021, discusses that iPLA2-VIA is necessary for the healthy aging of the neurons and muscle of the Drosophilae flies, since we found that the higher concentration of the gene is found in the early stages, this could indicate that the formation of neurological diseases start not later in life but on the early stages of the formation of the fly. This could be also expanded on the human being, since flies have a genome that is 60% homologous to that of humans, less redundant, and about 75% of the genes responsible for human diseases have homologs in flies (Ugur et al, 2016) we can assert that neurological diseases in humans could start to be expressed in early life stages such as in embryos and fetus, and the symptoms are only shown in later stages. The consistent expression of iPLA2-VIA in the larval and adult stages suggests that the gene plays a fundamental and possibly housekeeping role in the nervous system. This consistent expression may indicate that iPLA2-VIA is essential for basic neural functions such as membrane remodeling, signal transduction, and energy metabolism throughout development (Strokin *et al*, 2003; Ramanadham *et al*, 2015.

Moreover, the association of PLA2G6 mutations with neurodegenerative diseases in humans emphasizes the importance of understanding the functional roles of iPLA2-VIA in neural maintenance and repair mechanisms (Engel *et al*, 2010; Paisan-Ruiz *et al*, 2009). Our findings could inform future studies focused on the gene’s involvement in neural health and its potential as a therapeutic target for neurodegenerative diseases.

## 6 Conclusion

The consistent expression of iPLA2-VIA in the larval and adult stages suggests that the gene plays a fundamental and possibly housekeeping role in the nervous system. This consistent expression may indicate that iPLA2-VIA is essential for basic neural functions such as membrane remodeling, signal transduction, and energy metabolism throughout development. Moreover, the association of PLA2G6 mutations with neurodegenerative diseases in humans emphasizes the importance of understanding the functional roles of iPLA2-VIA in neural maintenance and repair mechanisms. Our findings could inform future studies focused on the gene’s involvement in neural health and its potential as a therapeutic target for neurodegenerative diseases. In conclusion, our study on the expression patterns of iPLA2-VIA in Drosophila nervous tissues revealed consistent expression in developmental stages, as expected by our initial expectations, with higher expression of Larvae than of Adult. These findings suggest fundamental roles for the gene throughout neural development and function. Future research involving more detailed analyses of RNA quality and alternative splicing could provide further insights into the regulatory mechanisms of iPLA2-VIA. Understanding these mechanisms may offer valuable information for studying similar processes in humans and developing strategies to combat neurodegenerative diseases, such as Parkinson’s and Alzheimer’s.

## Acknowledgments

We thank Yeshiva University for providing the labs and Dr. Josepha Steinhaeur for allowing the use of her lab and all the help.

## Appendix

Figure 1 Network Link:

https://www.ndexbio.org/#/network/cfd213da-2e99-11ef-9621-005056ae23aa?accesskey=9635018a0c489cac964d770d460e7160766f5980a4b03f9e5791e8e5319b4b93

Figure 2 RNA Ladde and Gel:

**Figure 2:**
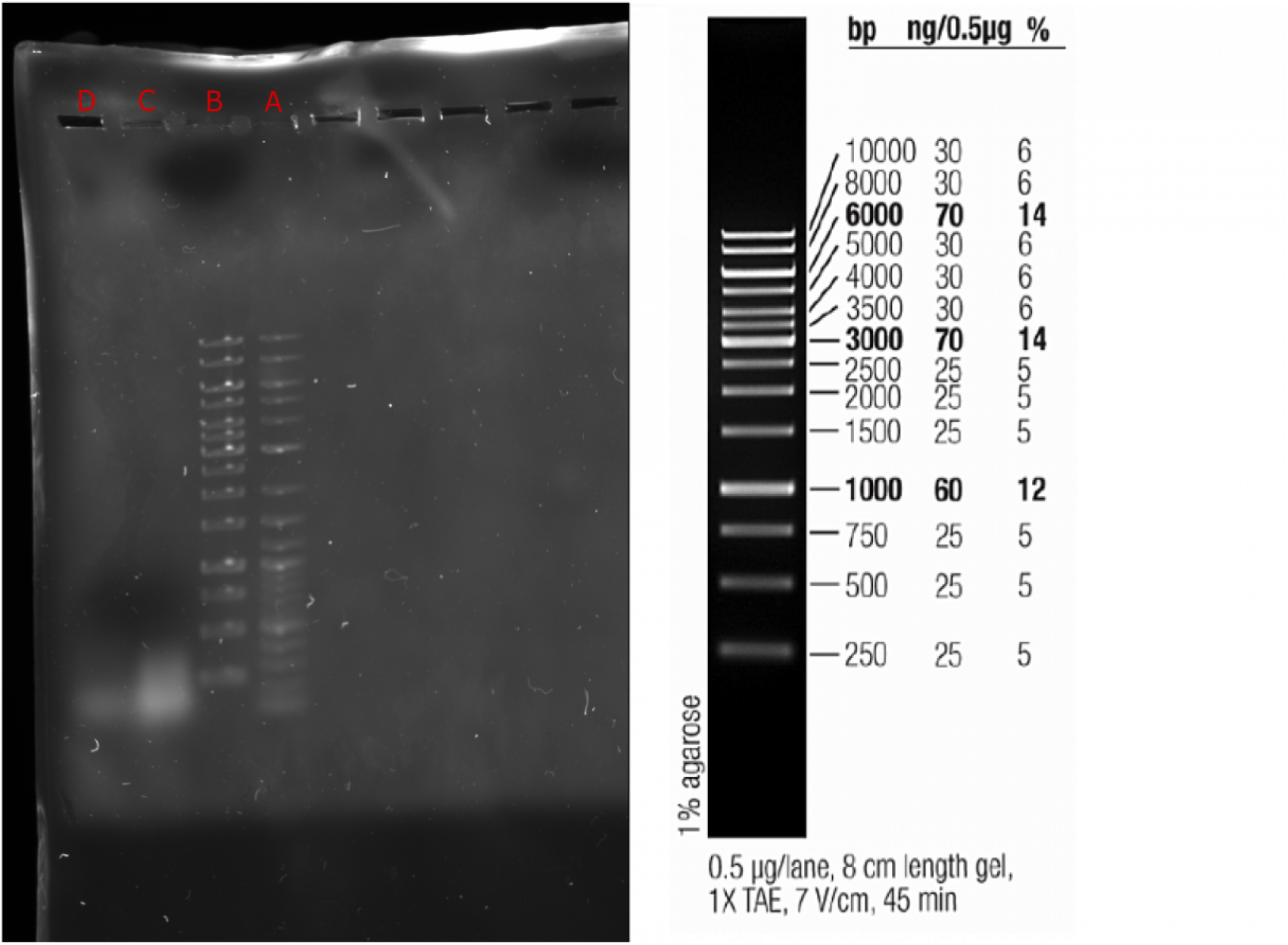
**Gel Image: D) stands for Adults; C) for Larvae; B) for 1kb Ladder, and A) for 100 kb Ladder. The expected position of the RNA for both Larvae and Adults can be seen close to the 500 bp marker, our results indicate that the RNA was close to the 250 bp marker. An RNA ladder on the right is provided for reference.**

## Notes

### Competing Interest Statement

The authors have declared no competing interest.

### Summary of Updates

bibliographic order for inline citation of references. The second figure in the manuscript was labeled as Figure 4. Removed citations in the conclusion and provided appropriate references.

